# A new *Paramoeba* Isolate from Florida Exhibits a Microtubule-Bound Endosymbiont Closely Associated with the Host Nucleus

**DOI:** 10.1101/2025.03.10.642444

**Authors:** Yonas I. Tekle, Atira R. Smith, Michael McGinnis, Saron Ghebezadik, Priyal Patel

## Abstract

The genera *Paramoeba* and *Neoparamoeba*, within the family Paramoebidae (order Dactylopodida), are distinguished by their dactylopodial pseudopodia and the presence of an intracellular eukaryotic symbiont, the *Perkinsela*-like organism (PLO). Taxonomic classification within these genera has been challenging due to overlapping morphological traits and close phylogenetic relationships. Most species are marine, with some acting as significant parasites, contributing to sea urchin mass mortality and serving as causative agents of Amoebic Gill Disease (AGD). Despite their ecological and economic importance, many aspects of their diversity, biology, evolution, and host interactions remain poorly understood. In this study, we describe a novel amoeba species, *Paramoeba daytoni* n. sp., isolated from Daytona Beach, Florida. Morphological and molecular analyses confirm its placement within the *Paramoeba* clade, closely related to *P. eilhardi, P. karteshi,* and *P. aparasomata*. Phylogenetic assessments using 18S and COI markers demonstrate the limitations of 18S gene for species delineation, highlighting COI as a more reliable genetic marker for this group. Additionally, observations on PLO morphology, movement, and microtubule association provide insights into the endosymbiotic relationship, reinforcing the need for further research into this unique eukaryote-eukaryote symbiosis.

---

The genera, *Paramoeba* and *Neoparamoeba*, comprise amoebae characterized by dactylopodial type of pseudopodia within the family Paramoebidae (order Dactylopodida). One of the identifying features of *Paramoeba*/*Neoparamoeba* has been the presence of an intracellular eukaryotic symbiont, known as the *Perkinsela*-like organism (PLO). In addition to this the general dactylopodia morphotype and micro-scale on cell surface are among characters used in the taxonomy of the group (Page 1987).

*Paramoeba/Neoparamoeba* species primarily inhabit marine environments, with some species known to play significant parasitic roles in marine organisms (Jones 1985, Dykova et al. 2005, Young et al. 2007, Sprague et al. 1969). For example, *Paramoeba invadens* has been implicated in mass mortalities of sea urchins in Nova Scotia (Feehan et al. 2013), while other *Paramoeba* spp. have been identified as causative agents of Amoebic Gill Disease (AGD) in Atlantic salmon (*Salmo salar*), a major issue for aquaculture industries (Foster and Percival 1988). Despite their ecological and economic significance, their diversity, evolution, life cycle and pathobiology remain poorly understood.

The relationship between *Paramoeba* and its PLO symbiont represents a unique and seemingly stable eukaryote-eukaryote endosymbiosis. Initially misinterpreted as an organelle or secondary nucleus (Perkins and Castagna 1971, Schaudinn 1896, Janicki 1912, Chatton 1953, Grell 1961). The PLO was later identified as a kinetoplastid endosymbiont closely related to *Ichthyobodo necator* (Dyková et al. 2003, Hollande 1981, Tanifuji et al. 2011). The PLO is invariably located near the host amoeba’s nucleus and appears to be obligatory for the host’s survival, as neither organism has been successfully cultured independently (Dyková et al. 2003). Despite the symbiosis, genomic analyses suggest that significant genetic integration between the host and PLO has not occurred, indicating a relatively recent adaptation to intracellular life (Tanifuji et al. 2011).

Molecular phylogenetic studies of *Paramoeba* and their PLO have relied primarily on molecular markers such as small subunit-rDNA (18S) gene, revealing a strong signal of co-evolution between the amoebae and their PLO endosymbionts (Caraguel et al. 2007, Dykova et al. 2008, Young et al. 2014, Sibbald et al. 2017). However, taxonomic classification within *Paramoeba/ Neoparamoeba* remains challenging due to limited phylogenetic signal in the 18S gene and the overlapping morphological features between *Paramoeba* and *Neoparamoeba*.

Historically, the presence or absence of microscales on the cell surface was used as a distinguishing feature, but recent molecular phylogenetic evidence suggests that micro-scale presence alone may not be a reliable taxonomic character (Feehan et al. 2013, Young et al. 2014). Additionally, the recent description of a *Paramoeba* isolate lacking a PLO and a closer reexamination of previously described species in the genus *Korotnevella*, further complicates the foundational taxonomic scheme used in the group (Volkova et al. 2019).

In this study, we present a novel amoeba isolate from Daytona Beach, Florida, that is closely related to a clade consisting of the type species, *Paramoeba eilhardi* Schaudinn, 1896, based on both morphological and molecular data. We provide a detailed analysis of its morphology, including observations on the association between the PLO and the host nucleus using light and immunocytochemistry techniques. Furthermore, we present a molecular phylogenetic analysis of the group, illustrating the placement of the new isolate and reinforcing the congruence between species phylogeny and PLO co-evolution.

## MATERIALS AND METHODS

### Cell culturing and DNA isolation

A *Paramoeba* species designated as YT-Flat isolate presented in this study was isolated from a mix of seaweed and sand samples collected from Daytona Beach Florida, USA (29.2108° N, 81.0228° W) in the year 2022. A mix of the collected samples was cultured in a peri dish with 3% artificial seawater solution with autoclaved grains of rice (a food source for bacteria growing with the amoebae). After several rounds of culturing the isolate appeared in large numbers in one subculture and was subsequently isolated to establish a monoclonal culture. While several approaches were used to establish monoclonal cultures, a tiny flagellate persistently appears in our subcultures. The amoeba was observed to engulf these tiny flagellates as alternative food source to bacteria that came from nature with the original samples.

### Light microscopy and Immunocytochemistry

Observations on the general morphology of amoeba cells grown in plastic Petri dish cultures and glass slides were recorded using ECLIPSE Ti2 (Nikon Corporation, Japan). Using various functionalities in the NIS-Elements software (Nikon Corporation) we collected still images and timelapse videos with phase contrast and Differential Interference Contrast (DIC) settings.

Measurements were taken using the NIS software. Staining of subcellular structures including plasma membrane, cytoskeleton (microtubule) and DNA were studied using immunocytochemistry (ICC) methods as described in Tekle and Williams (2016).

### DNA extraction and PCR amplification

Clean cultures of YT-Flat isolates were used to extract total genomic DNA using Quick-DNA ™ MicroPrep DNA Extraction Kit (Zymo Research, Irvine, California, cat. no. D3020) per manufacturer’s instructions. Primers for SSU-rDNA (18S) gene were from Medlin et al. (1988). Phusion DNA polymerase, a strict proofreading enzyme, was used to amplify the genes of interest with PCR conditions described in Tekle et al. (2013). Amplicons from multiple PCR reactions were purified using QIAquick PCR Purification Kit (cat.no. 28106) and pooled together to get enough concentration for Oxford Nanopore sequencing. The cytochrome C oxidase subunit 1 (COI) gene was obtained from genome data of the isolate sequenced in Oxford Nanopore using whole genome amplification (WGA) of single cells as in Tekle et al. (2021).

### Nanopore sequencing and bioinformatics

PCR amplicons were quantified using a Qubit assay with the dsDNA broad range kit (Life technologies, Carlsbad, CA, USA). Library preparation was performed using the Oxford Nanopore Rapid Barcoding Kit (SQK-RBK114.24) according to the manufacturer’s protocol. A total concentration ranging from 100–200 ng of input DNA was used for barcoding. Barcoded adapters were ligated directly to the DNA via a transposase reaction, which simultaneously fragmented the DNA. Following adapter ligation, barcoded samples were pooled and purified using AMPure XP beads to remove excess adapters and short fragments. The final library concentration was determined using a Qubit fluorometer before loading onto the flow cell. A R10.4.1 flow cell was primed with sequencing buffer and loading mix as per the Oxford Nanopore protocol. The prepared library was then loaded onto the flow cell, and sequencing was initiated using the MinKNOW software (MKE_1013_v1_revDJ_16Jan2025) for a 48 hrs. run.

Raw sequencing reads were base called and demultiplexed using Guppy basecalling software. Quality control of the reads was performed using FastQC. Initial analysis of the nanopore reads were conducted using NCBI Blast. To account for errors in sequencing such as indels of the raw nanopore reads, we used a pipeline amplicon_ sorter v2022-03-28 (Vierstraete and Braeckman 2022) before using them for downstream analysis. The pipeline sorts and clusters nanopore reads based on length and sequence similarity, generating high-quality consensus metabarcodes. Amplicon_sorter considers all possible clusters when generating consensus sequences, making it well-suited for DNA metabarcoding analysis. Importantly, amplicon_sorter corrects indel errors during consensus calling, ensuring high-quality consensus sequences. We applied a conservative clustering approach, requiring sequences to be at least 97% similar to be grouped within a species cluster (--similar_species) and 98% similarity to merge consensus sequences (--similar_consensus). The length filter was set to a minimum length of 300 bps, and we performed three rounds of random sampling (--maxreads) to enhance the likelihood of capturing rare reads. Each cluster’s sequences were mapped back to their consensus using minimap2 v2.24, and final consensus sequences were polished with medaka v1.7.2 using the r103_sup_g507 model (https://github.com/nanoporetech/medaka).

### Phylogenetic, Pairwise and Approximately Unbiased (AU) analyses

Molecular phylogeny was reconstructed based on alignments of two markers (18S and COI) built using AliView (Larsson 2014) and MAFFT (Katoh and Standley 2013). We used four 18S gene datasets differing in the alignment methods used and number of characters retained. In order to retain maximum number of sites to resolve low diverging lineages, we prepared two pairs of 18S gene datasets with maximum number of characters (∼2300 bps) and with reduced data (ambiguous sites removed, ∼1750 bps) aligned using MAFFT (L-INS-i progressive method with --maxiterate 1000) and AliView (default setting). All alignments were carefully checked visually before further analysis. The final 18S gene alignment consisted of 166 ingroup and 10 outgroup taxa. For COI gene we used 19 ingroup and 4 outgroup taxa with a total number of 648 nucleotide sites. We also conducted a combined (2699 nucleotide sites) analysis of 18S and COI genes with 9 ingroup and 2 outgroup taxa. All pairwise comparison was made in MEGA (Kumar et al. 2001) using the K2P model.

Phylogenetic trees were reconstructed using the maximum likelihood algorithm in IQ-TREE (Nguyen et al. 2015, Hoang et al. 2018, Kalyaanamoorthy et al. 2017) and RaxML v.8.2.10 (Stamatakis 2014). For IQ-TREE the best-fit evolutionary model was determined automatically using the ModelFinder feature (-m AUTO) for all datasets including the combined COI and 18S genes (Kalyaanamoorthy, et al. 2017). Bootstrap support values (Hoang, et al. 2018) were estimated with 1,000 ultrafast bootstrap replicates (-B 1000) to assess node stability. RaxML was run with 100 initial parallel searches based on independent starting trees and nonparametric bootstrapping of the best tree with 1000 pseudo-replicates. The GTRGAMMAI model with default parameters was used. Bayesian inference was performed using MrBayes v.

3.2.6 (Ronquist et al. 2012) with GTR model and gamma model for among-site rate variation (4 rate categories) including proportion of invariable sites. Markov chain Monte Carlo analysis was performed in two runs of four simultaneous chains sampled for 2,000,000 generations for 18S gene datasets; 25% of the sampled trees were discarded as a burnin. The resulting phylogenetic trees were visualized and annotated using FigTree v1.4.4 (Rambaut 2018).

We used Approximately Unbiased (AU) tests (Shimodaira 2002) to test alternate tree topologies within the *Paramoeba/Neoparamoeba* phylogeny. Several loosely constrained topologies were optimized under MODEL in IQ-TREE. These optimized trees were compared with the main phylogeny, inferred using the AliView alignment with 2319 aligned nucleotide positions, using AU test with 10,000 RELL bootstrap replicates. The hypotheses that had p-AU ≥ 0.05 within the 95% confidence interval could not be rejected.

## RESULTS

### Morphology of *Paramoeba daytoni* n. sp

The newly isolated amoeba exhibits the typical morphology of the *Paramoeba* genus, characterized by several dactylopodial subpseudopodia that extend actively in locomoting and floating cells (Figure 1). During active movement, the amoeba predominantly adopts a flattened, triangular shape. Other observed forms include elongated or irregular shapes, particularly when changing direction or in transitional states (Data S1 timelapse). Subpseudopodia are primarily located in the mid-anterior region of the cell body, with occasional small projections visible throughout, including at the tapered posterior end (Figure 1F). Their number and position vary in locomoting cells, with an average of 7–10 subpseudopodia per cell (Figure 1). These projections can reach up to half the amoeba’s body length, measuring 3.55–31.34 µm (average 9.81 µm, N=33). During movement, subpseudopodia mostly extend laterally in the direction of movement, though some may flail dorsally (Figure 1E). Locomotion follows some degree of zigzag pattern, with leading subpseudopodia extending and new ones forming dynamically. Amoeba cells move at an average speed of 15.7 µm/min (range: 13–19.69 µm/min) at room temperature in plastic petri dishes. In the floating form, the amoeba exhibits multiple long, radiating subpseudopodia projecting from the central cell mass (Figure 1A). Cell measurements, including length and breadth, are 31.15–78.43 µm (average 54.73 µm, N=76) and 9.23–54.53 µm (average 23.85 µm, N=71), respectively. The nucleus, as observed in DIC images, averages 7.75 µm in diameter (range: 5.66–9.99 µm, N=21) (Figure 2). Most amoeba cells contain more than one PLOs, with some harboring up to four (Figure 2C). PLOs typically measure 2.02–4.67 µm in length (average 3.07 µm, N=44) and 1.76–3.19 µm in breadth (average 2.53 µm, N=44) . They are generally bean-shaped but may appear round depending on orientation (Figure 2). Further details on PLO association with the host nucleus and their intracellular movement are provided below.

**Figure 1.**
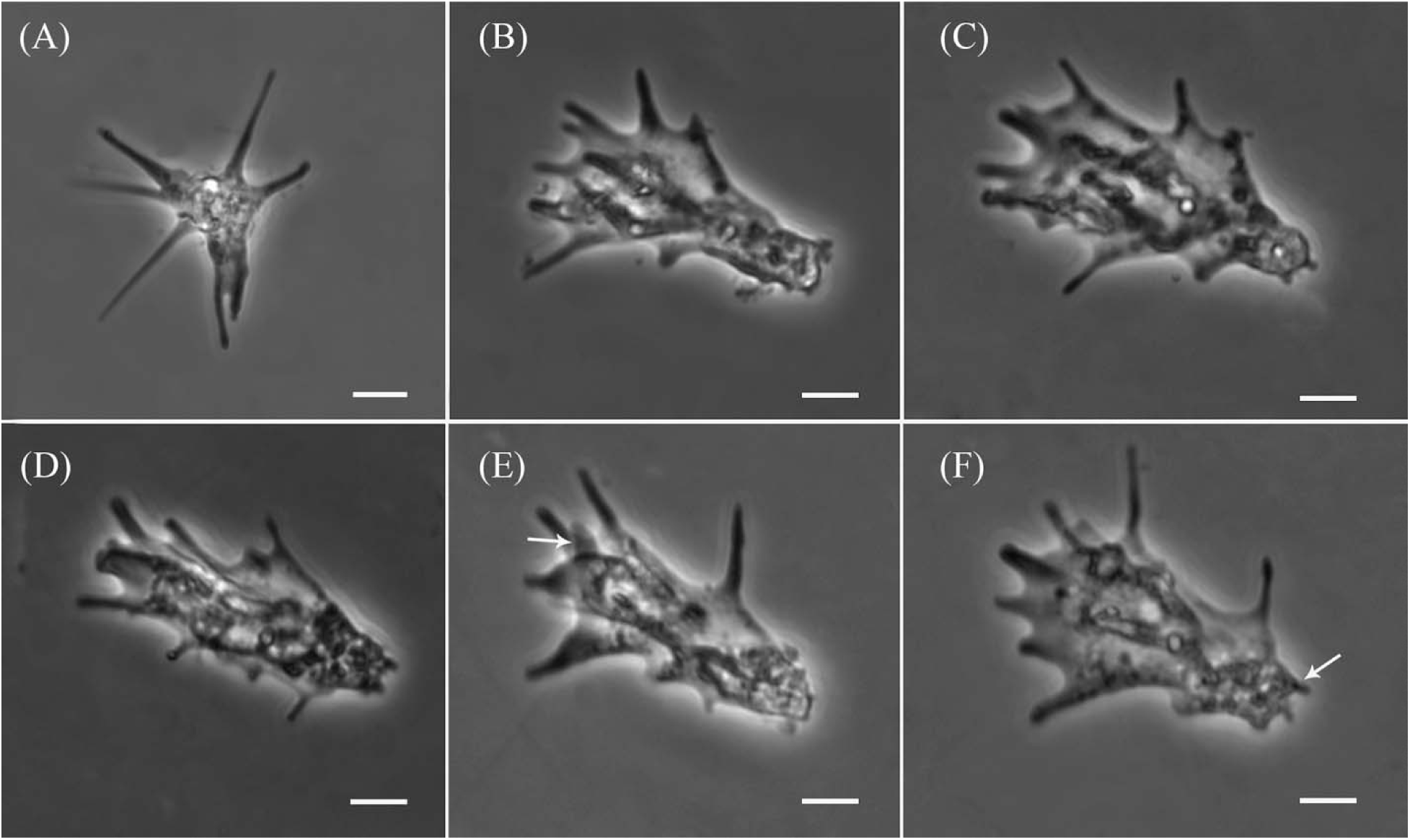
Light microscope phase-contrast micrographs of *Paramoeba daytoni* n. sp., showing its transition from a floating to fully locomotive forms. (A) Floating form. (B–F) Locomoting morphotypes, with long subpseudopodia in the anterior and mid-regions, upward-positioned flailing subpseudopodia (arrow, E) and shorter subpseudopodia at the posterior (arrow, F). All scale bars represent 10 µm.

**Figure 2.**
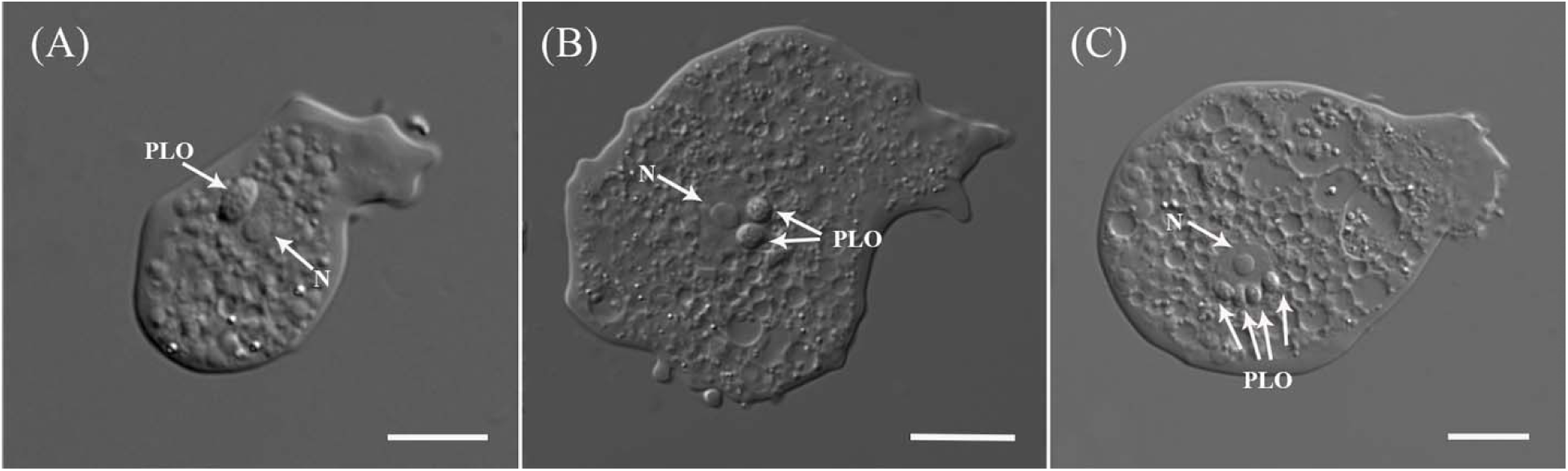
DIC micrographs of *Paramoeba daytoni* n. sp. showing the nucleus (N) and variation in the number of *Perkinsela*-like organism (PLO). (A) A single oval-shaped PLO. (B) Two round-shaped PLOs. (C) An isolate with the maximum recorded number of PLOs (4) for the new species. All scale bars represent 10 µm.

### Immunocytochemistry staining and PLO movement observation

We used both immunocytochemistry and DIC imaging timelapse videos to observe the amoeba and its association with its endosymbiont (PLO) in the new isolate (Figures 3, S1, Data S1 timelapse). Immunocytochemistry staining provided a general outline of the cell using CellMask (plasma membrane), DAPI (nucleus), and microtubule (MT) cytoskeleton markers (Figures 3, S1). The overall microtubule architecture of the new isolate resembles that of most amoebae, consisting of a complex network distributed throughout the cell body, sometimes extending into the subpseudopodia (Figures 3C, S1). Although our staining technique did not preserve subpseudopodia well, some staining shows that the extending subpseudopodia are supported by a few microtubules (Figure S1D). Around the host nucleus, a network of MTs is observed to provide structural support (Figures 3C, S1C).

**Figure 3.**
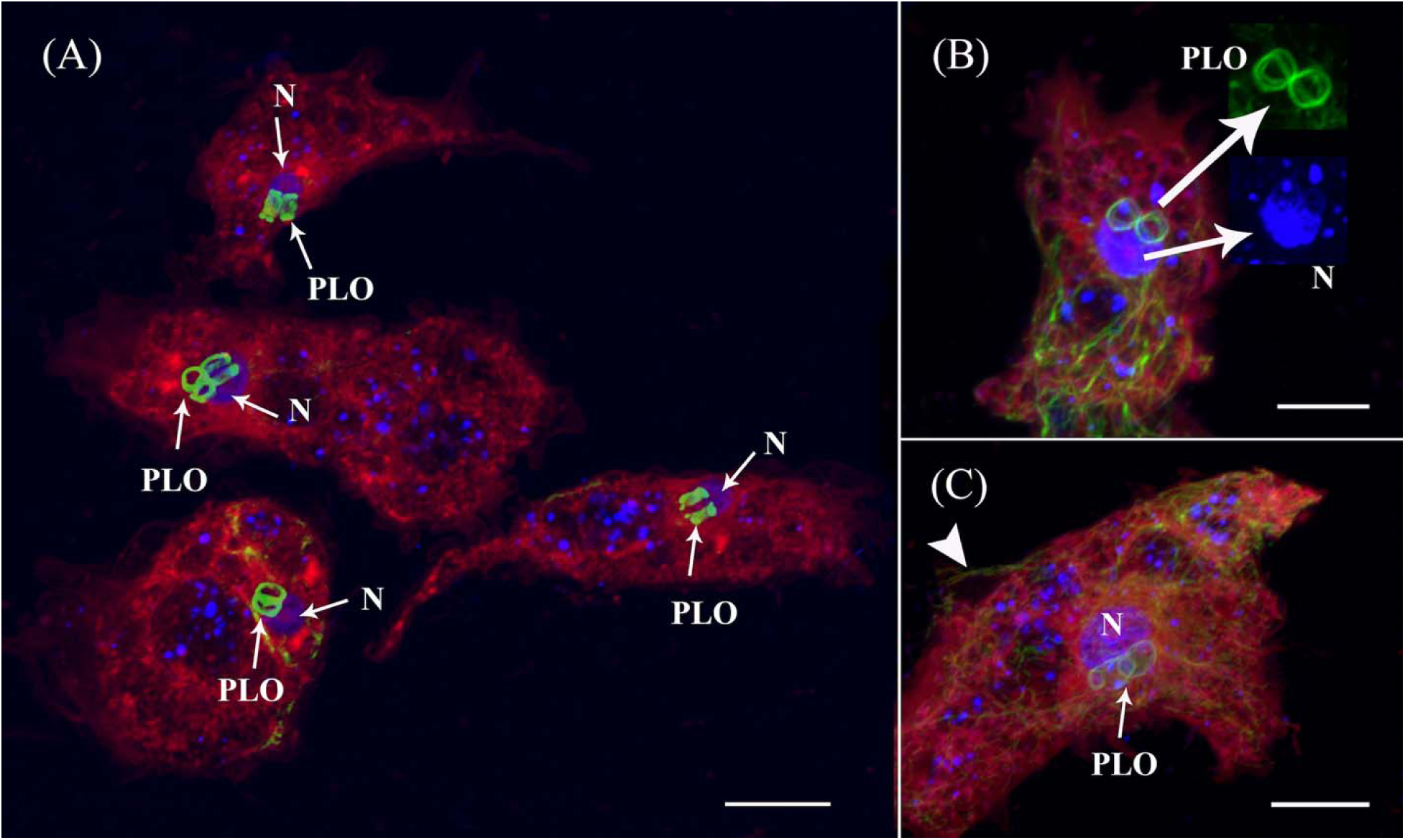
Confocal maximum intensity projection images of microtubules (green), DNA (blue), and the plasma membrane (red) in *Paramoeba daytoni* n. sp. (A) Four amoeba cells with varying numbers of PLOs. Strongly stained interconnected microtubules (MTs) rings of PLO are observed adjacent to the nucleus (N, arrows). The host amoeba MTs are not well preserved in these specimens. (B) An amoeba cell with two PLOs forming a figure-eight shape, consisting of more than one microtubule bundle. Insets (arrows) provide close-up images of the PLO microtubules and the host amoeba nucleus (N). Note in the inset of nucleus that part of the nucleus staining is overshadowed by the overlaying PLOs. The plasma membrane and PLO nuclei are not distinctly visible in any of our staining, aside from randomly dispersed DAPI-stained small dots, which are likely artifacts or mitochondrial DNA (A–C). (C) An amoeba cell with bean-shaped, overlapping PLO microtubules. Note the microtubule extension over the side subpseudopodia (arrowhead).

All analyzed PLOs are tightly enwrapped by MTs, which stain more intensely than any other host cell MTs (Figures 3, S1). One or more MT bundles are observed encircling each PLO, often forming a figure-eight shape when two PLOs are present (Figure 3B) or connecting tangentially when more than two are observed (Figure 3A). Occasionally, PLOs appear stacked or partially overlapping but are never fully separated (Figure 3). While some staining suggests a possible connection between host MTs around the nucleus and PLO-associated MTs (Figure S1C), the PLO-associated MTs generally appear distinct from the host microtubular network in both intensity and organization.

The host nucleus stains strongly with DAPI, and in some cells, variations in ploidy are evident based on nuclear size and nucleolar number (Figure S1A). However, both CellMask and DAPI staining fail to reveal a plasma membrane or DNA signal in the PLO, despite multiple staining attempts. Whether this is due to fixation issues or other factors remains unclear. The presence of PLOs in our ICC staining is confirmed solely by their strong MT staining (Figures 3, S1).

To further investigate PLO dynamics, we analyzed live-cell DIC timelapse videos to explore its association with the host nucleus (Data S1 timelapse). In all observations, the PLO remains closely associated with the nucleus. During amoeba locomotion or cytoplasmic streaming, both the nucleus and PLO move together through the cytoplasm. The PLO appears to move around the circumference of the nucleus but never completes a full rotation (Data S1 timelapse). Instead, its movement seems restricted, oscillating back and forth along most of the nucleus’s periphery, suggesting the presence of a constraining force.

### Molecular data collection via Oxford nanopore sequencing

We used Nanopore sequencing, as part of a metabarcoding and genome sequencing project, to obtained genetic data for this study. Using universal primers, we successfully amplified the full-length 18S gene of the host. When this amplicon was sequenced with Oxford Nanopore, our raw data contained multiple sequences representing the host 18S gene, along with a few co-amplified endosymbiont (PLO) ribosomal DNA sequences. The host 18S gene sequences were polished using the Amplicon-sorter pipeline, resulting in high-quality sequences. However, due to the low representation of PLO 18S gene sequences, it was not possible to reconstruct a reliable consensus sequence using the same pipeline. The PLO 18S gene sequences contained indels and regions with low alignment confidence due to potential sequencing errors. A preliminary manually curated alignment and phylogenetic analysis of the PLO 18S gene sequences showed a congruent grouping of the new isolate with the host 18S gene phylogeny (see below). PCR amplification of the COI gene using universal primers was unsuccessful. However, COI gene data for the new isolate was obtained from the genome sequencing project, where multiple copies of the gene were found as part of the mitochondrial genome.

### Sequence divergence and phylogenetic analyses

Given the variation observed in 18S gene phylogenies in our preliminary analyses, along with the high sequence representation and complex intra- and interspecific divergences reported in previous studies (Feehan et al. 2013, Volkova et al. 2019), we employed multiple approaches to analyze both markers (18S and COI). These approaches included pairwise comparisons within and among various clades, evaluating alignments with different character sets and alignment methods, and using different algorithms to reconstruct phylogenies.

The maximum intrastrain variation of 18S gene in *Paramoeba/Neoparamoeba* species ranges from 1.33% to 5.54% (Table S1), with the lowest observed in *Paramoeba longipodia* (1.33%) and the highest in *Paramoeba aestuarina* (5.72%). Notably, the maximum intrastrain variation and the minimum interspecific variation in *Paramoeba/Neoparamoeba* species show clear overlap (Figure 4, Table S2), meaning that some intrastrain variations are as high as the interspecific variations recorded in certain groups. While 18S gene generally exhibits low variation at both intra- and interspecific levels, COI gene shows lower intrastrain variation but significantly higher interspecific divergence across all analyzed species. The interspecific COI gene divergence consistently exceeds 12%, well above the threshold used for species delimitation in amoebae and other organism. The new isolated exhibits COI gene sequence divergences of 14.57%, 15.91%, and 13.48% compared to *P. eilhardi*, *P. aparasomata*, and *Paramoeba karteshi*, respectively.

**Figure 4.**
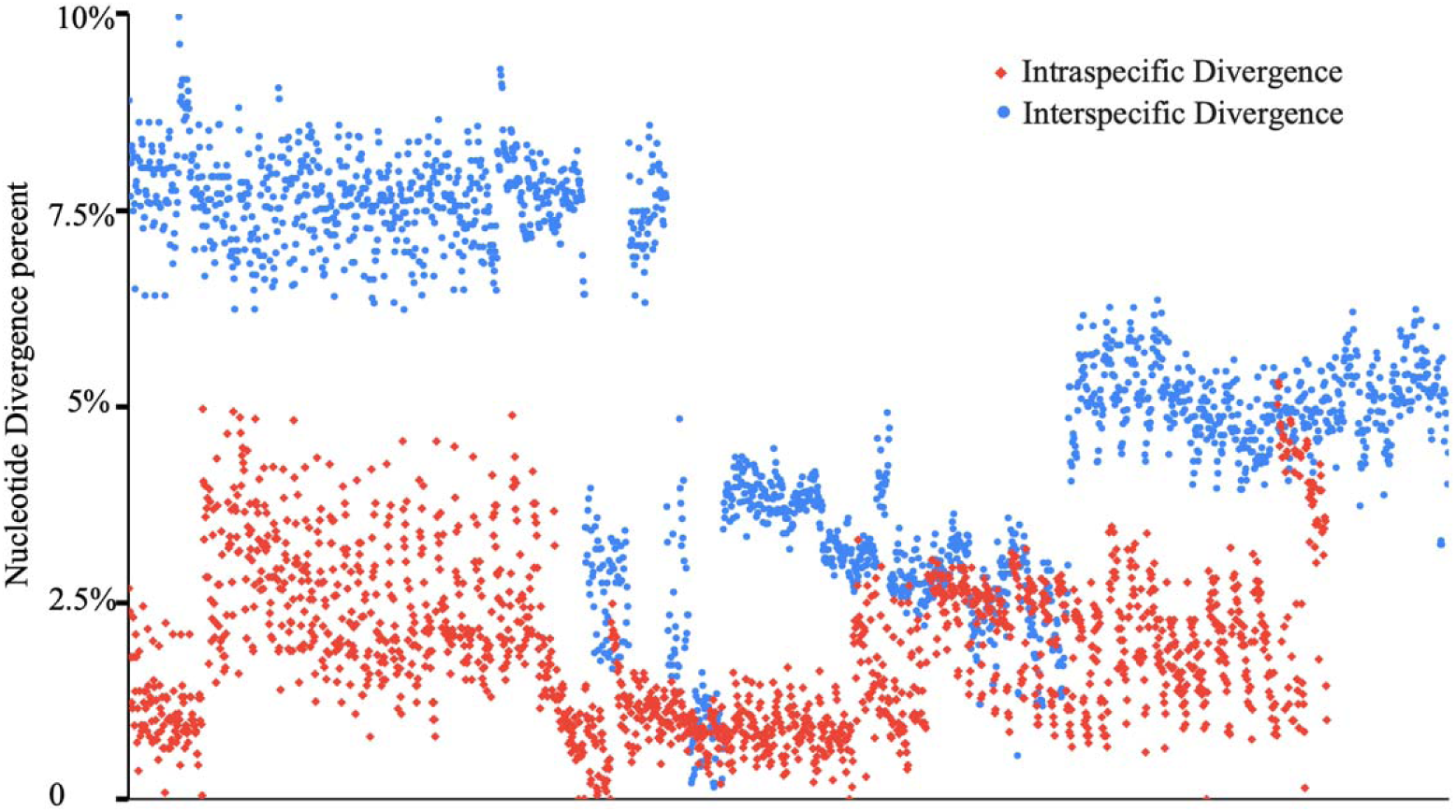
Comparison of *Paramoeba/Neoparamoeba* clade species based on pairwise 18S gene divergence, illustrating overlapping intrastrain and interspecific percentage values.

The 18S gene phylogenetic tree, reconstructed using different alignments, character sets, and three tree reconstruction algorithms, mostly yielded consistent results (Table S3). The core clades recovered with robust support in all analyses include (*P. invadens* + *P. branchiphila*) + *P. aestuarina* (IBL clade, species initials used for short representation in clade names) and *P. pemaquidensis* + *P. aestuarina* (PA clade) (Figure 5, Table S3). The new isolate consistently groups within a clade that includes *P. eilhardi*, *P. aparasomata*, and *P. karteshi* (Figure 5).

**Figure 5.**
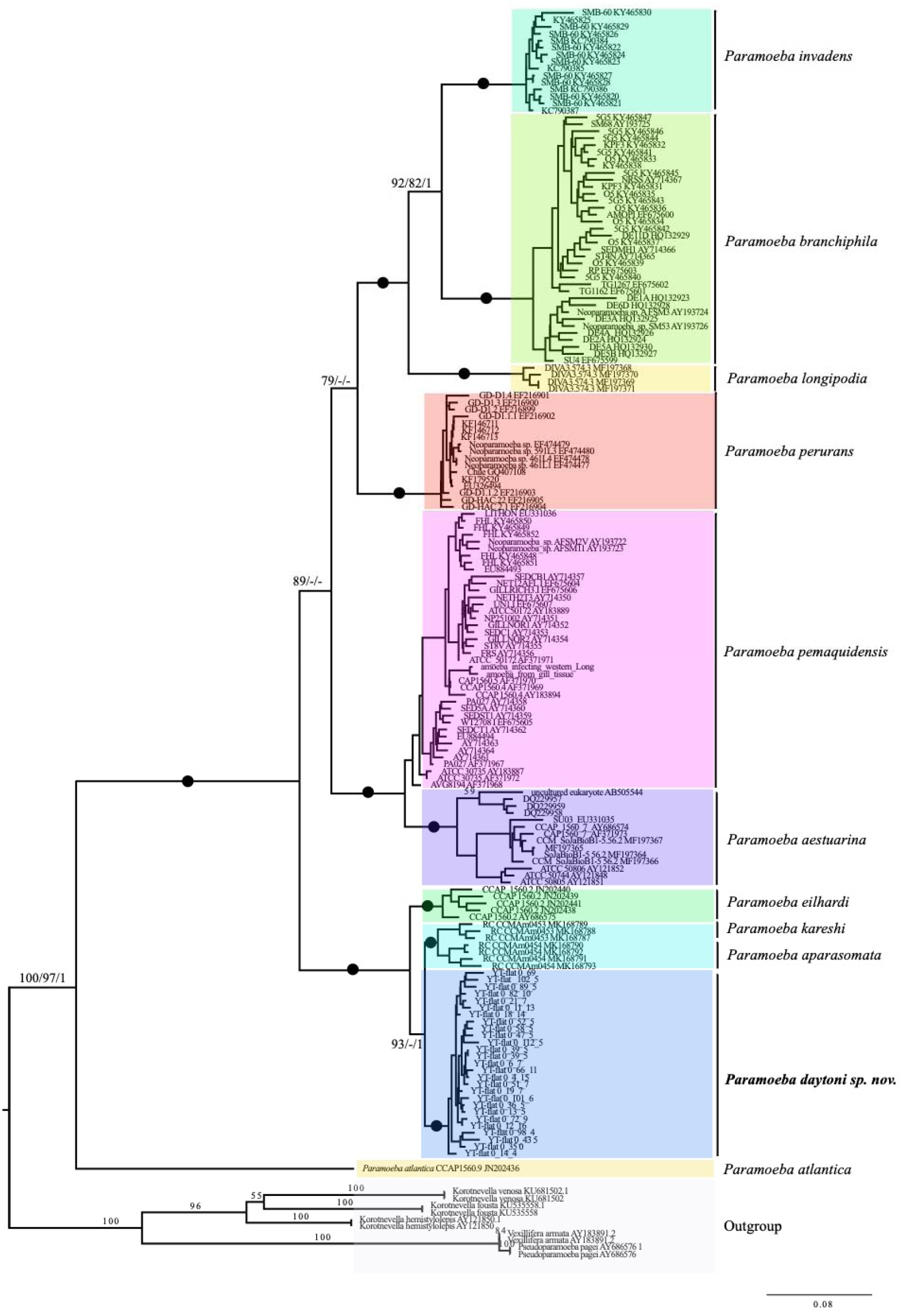
IQ-TREE maximum likelihood phylogeny of 18S gene based on 167 ingroup taxa of *Paramoeba/Neoparamoeba*, highlighting the position of *Paramoeba daytoni* n. sp. The alignment, generated using AliView, includes 2,319 nucleotide positions. Numbers at nodes represent IQ-TREE/RAxML bootstrap values and Bayesian posterior probabilities, respectively. Solid black circles indicate full support across all analyses. Branches are drawn to scale.

However, in 50% of the analyses, it clusters either with *P. eilhardi* or with *P. aparasomata* + *P. karteshi* (Table S3). The support and placement of the new isolate vary depending on the algorithm and the number of characters analyzed (Table S3).

Most analyses place the new isolate with *P. karteshi* and *P. aparasomata* (E-DAK clade) when more sites are included. However, when using a more conservative alignment with fewer sites, it clusters with *P. eilhardi* (ED-AK clade) (Table S3). Given the low interspecific divergence, in *Paramoeba/Neoparamoeba* species, ranges, we present a phylogeny based on a larger dataset (2319 sites) in this study (Figure 5). *P. atlantica* is consistently recovered as a basal lineage in all analyses (Figure 5, Table S3). However, depending on the dataset and methodology used, we observed varying topologies regarding the placement of *P. perurans* and the EDAK clade in relation to the IBL and PA clades (Table S3).

We conducted an AU test to assess the robustness of the various topologies (Table S4). The grouping of the new isolate with *P. aparasomata* + *P. karteshi* received a higher probability than its grouping with *P. eilhardi*. Although the latter sister-group relationship cannot be rejected, the COI gene phylogeny and the combined 18S and COI genes analysis favor the sister relationship of our new isolate with *P. aparasomata* + *P. karteshi*, consistent with the AU test results (Figures 6, S2). All alternative topologies that place *P. perurans* (PE) with the EDAK+PA clade receive higher probability than those that position EDAK or PE at the base of the tree, branching before *P. atlantica* (Figure 5, Table S4).

**Figure 6.**
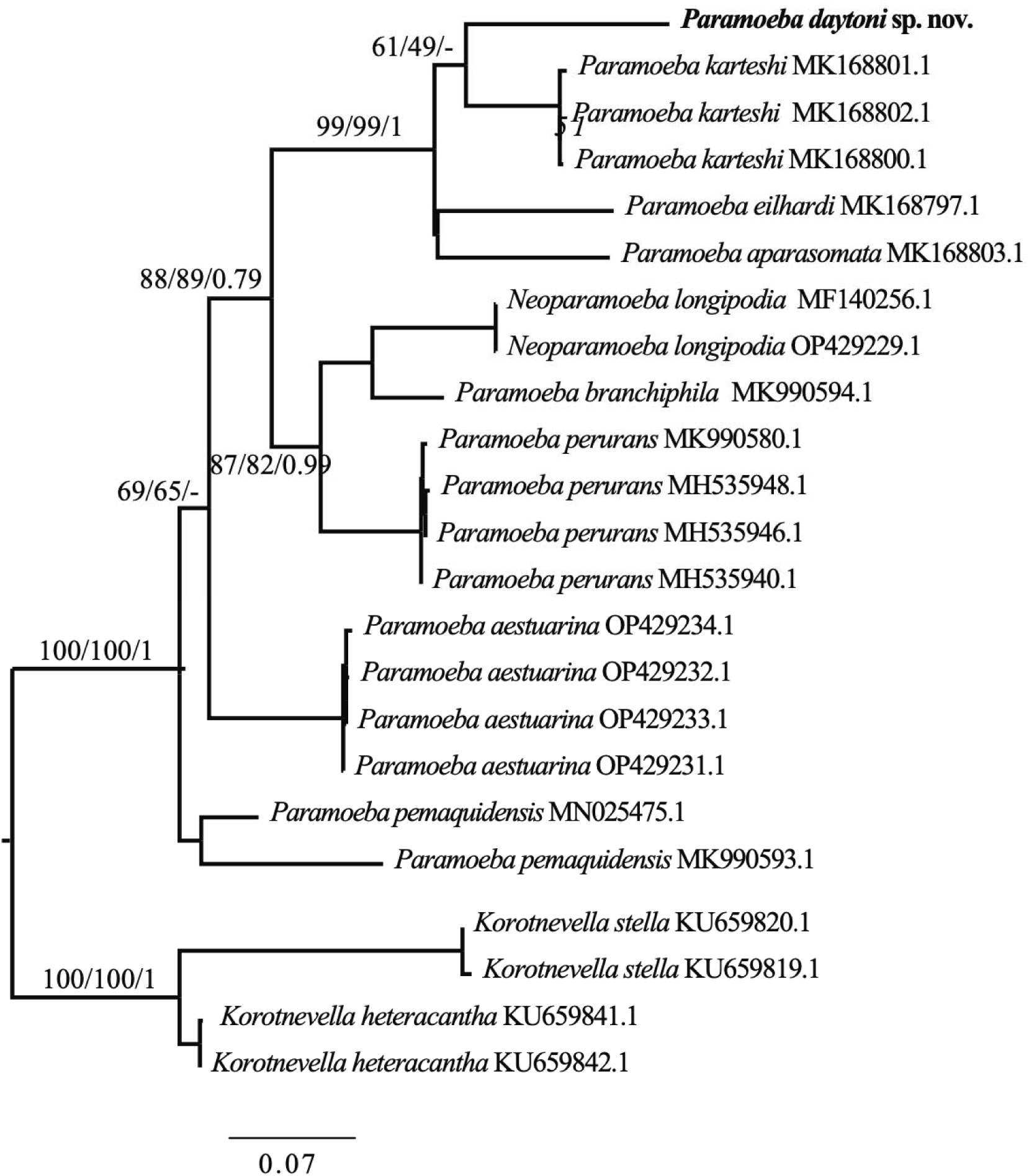
IQ-TREE maximum likelihood phylogeny of the COI gene based on 19 ingroup taxa of *Paramoeba/Neoparamoeba*, highlighting the position of *Paramoeba daytoni* n. sp. The alignment, generated using AliView, includes 648 nucleotide positions. Numbers at nodes represent IQ-TREE/RAxML bootstrap values and Bayesian posterior probabilities, respectively. Solid black circles indicate full support across all analyses. Branches are drawn to scale.

In the COI gene phylogeny, the new isolate robustly clusters within the EDAK clade (Figure 6). It forms a sister group with *P. karteshi*, while *P. eilhardi* and *P. aparasomata* cluster together; however, both relationships receive little to no support (Figure 6). Overall, the internal relationships within this clade are weakly supported, with the placement of *P. perurans* alongside *P. longipodia* and *P. branchiphila* receiving only moderate support (Fig. 6). This clade also forms a sister group with the EDAK clade with moderate support, whereas *P. pemaquidensis* and *P. aestuarina* do not cluster together as seen in the 18S gene phylogeny.

The concatenated analysis of COI and 18S genes yielded results similar to those of the individual COI and 18S phylogenies (Figure S2). Unlike in the COI phylogeny, *P. pemaquidensis* and *P. aestuarina* form a strongly supported sister group. The new isolate clusters as a sister group to *P. karteshi* and *P. aparasomata*, though with weak support (Figure S2). The remaining topology largely mirrors that of the COI gene phylogeny (Figure 6).

We also attempted to reconstruct a PLO phylogeny using the unpolished 18S gene sequences generated in this study along with all available PLO 18S gene sequences from NCBI. Surprisingly, despite the presence of indels and potential sequencing errors in our isolate, it grouped robustly with the PLO sequences of *P. karteshi* and *P. eilhardi* (Figure S3). The placement of the new isolate outside of the *P. karteshi* + *P. eilhardi* cluster may be due to taxon sampling limitations (as no PLO sequences for *P. aparasomata* are available) or potential sequencing errors. Overall, the topology of the PLO 18S gene phylogeny is largely congruent with the phylogenies generated in this (Figure 5) and a previous study (Volkova et al. 2019).

## DISCUSSION

A new species from Florida coastal lines, *Paramoeba daytoni* n. sp.

During an excursion to the northeastern coastal regions of Florida (Atlantic Ocean), we discovered a diverse range of amoeboids, including the newly isolated species. Our findings are consistent with previous reports indicating that the Florida coastline harbors a rich and largely unexplored diversity of microbial amoeboids (Rogerson and Gwaltney 2000, Davis et al. 1978, Sawyer 1980).

The new isolate is a free-living amoeba obtained from samples collected at a public beach in Daytona Beach. The *Paramoeba/Neoparamoeba* clade include both free-living and parasitic marine species (Volkova et al. 2019, Foster and Percival 1988, Feehan et al. 2013, Young et al. 2007, Jones 1985). All morphological and molecular data support its classification within the *Paramoeba/Neoparamoeba* clade (Figures 1-6). The high morphological plasticity among described species of *Paramoeba/Neoparamoeba*, has led to taxonomic confusion within the group (Volkova et al. 2025, Dyková et al. 2000). This indicates a high probability of cryptic species diversity within the group. In addition to general morphology and ultrastructural characters, the presence and number of PLO, within the *Paramoeba/Neoparamoeba* clade has been used as diagnostic feature (Page 1987, Hollande 1980). However, inclusion of lineages that lack the PLO within the genus highlights the limitations of this feature as a taxonomic marker (Volkova et al. 2019). While general morphology is useful for genus-level identification, species-level classification must be complemented with molecular data.

The newly described species, *Paramoeba daytoni* n. sp., clusters robustly within a clade that includes *P. eilhardi*, *P. aparasomata*, and *P. karteshi* (Figures 5, 6). Based on published morphometric data from Volkova et al. (2019), *P. daytoni* n. sp., is slightly larger in length and has the highest recorded number of PLO endosymbionts (up to 4) within the EDAK clade. The PLO diameter of *P. daytoni* n. sp. (∼3 µm) is more similar to that of *P. karteshi* than *P. eilhardi*, which has the largest recorded PLO size, 7 µm (Volkova et al. 2019). Similarly, the nucleus diameter of *P. daytoni* n. sp. (∼7 µm) is closer to that of *P. karteshi* and *P. aparasomata* rather than *P. eilhardi* (∼8 µm). These morphological characteristics, along with molecular phylogenetic analysis and sequence divergence of the barcode marker, COI, support the description of our isolate as a novel species.

### Challenges in the molecular Phylogeny of *Paramoeba*/*Neoparamoeba*

A substantial amount of molecular data (over 160 18S gene sequences) representing *Paramoeba* and *Neoparamoeba* strains is available in public databases. Phylogenetic analysis based on 18S gene has been useful in identifying clusters of both named and unnamed *Paramoeba*/*Neoparamoeba* species. However, previous studies have shown that the distinction between the two genera is blurred in 18S gene phylogenies (Volkova et al. 2019, Volkova and Kudryavtsev 2017, Sibbald et al. 2017). Based on these findings, some authors have suggested synonymizing *Neoparamoeba* with *Paramoeba* (Feehan et al. 2013).

A major challenge in 18S gene phylogeny is the high degree of sequence similarity (low interspecific divergence) among named *Paramoeba/Neoparamoeba* species (Table S2). Our analysis reveals an overlap between intrastrain and interspecific divergence among published 18S sequences of *Paramoeba*/*Neoparamoeba*, complicating species delimitation and affecting the resolution of the phylogenetic tree (Figure 4). The limitations of the 18S gene in Amoebozoa phylogenetics are twofold: while some amoebozoan taxa exhibit low interspecific variation, others display staggeringly high intragenomic variation in the 18S gene (Kudryavtsev and Gladkikh 2017, Zlatogursky et al. 2016). Additionally, in certain amoebozoans, 18S gene sequences exhibit such high degree of divergence that universal primers fail to amplify them (Tekle et al. 2008). These challenges the universal application of 18S gene for phylogenetic reconstruction and species delineation in some groups of the Amoebozoa. To address these limitations, comparative phylogenomic analyses using large-scale transcriptomic data have been employed to identify cryptic species (Tekle and Wood 2018) and resolve the phylogenetic placement of enigmatic taxa (Kang et al. 2017, Tekle et al. 2022). Additionally, the use of the COI mitochondrial gene as a DNA barcode marker has proven to be a practical solution for species delimitations in several amoebozoan lineages (Tekle 2014, Nassonova et al. 2010).

For *Paramoeba*/*Neoparamoeba*, COI gene sequence data is growing, with nine COI gene sequences currently available for described or named species. Using these data, we constructed phylogenies and performed pairwise comparisons to gain insights into the evolution of these species. Unlike 18S gene, the interspecific divergence observed in *Paramoeba*/*Neoparamoeba* based on COI gene is significantly higher (>12%), well above the threshold typically used for species delimitation using this marker (Tekle 2014). This is surprising given the low and overlapping intra-interspecific divergence observed in the 18S gene (Figure 4, Tables S2,3). For example, in *Cochliopodium*, which also exhibits low interspecific divergence in 18S gene, the COI gene divergence is also very low and close to the 2% threshold commonly used for species demarcation (Tekle 2014). A similar trend has been reported in other amoebae (Nassonova et al. 2010). These findings suggest that COI gene is a more reliable marker for species delineation within the *Paramoeba*/*Neoparamoeba* clade. A recent preprint examining the intraspecific variability, including ITS (Internal Transcribed Spacer) genes, of *Neoparamoeba pemaquidensis* supports this conclusion, further reinforcing the expectation that significant cryptic diversity exists within the *Paramoeba*/*Neoparamoeba* clade.

The phylogenetic relationships within the *Paramoeba/Neoparamoeba* clade, based on 18S and COI markers, reveal several consistently recovered clades (Figures 5, 6, S2). These include the IBL clade (*P. invadens*, *P. branchiphila*, and *P. aestuarina*) and the PA clade (*P. pemaquidensis* and *P. aestuarina*). *Paramoeba atlantica*, which has a highly divergent 18S gene sequence compared to other *Paramoeba/Neoparamoeba* species, consistently groups as the most basal lineage (Figure 5, Table S3). However, the positions of *P. perurans* and the EDAK clade vary between analyses and generally receive weak support (Figure 5, Table S3). Their placement within the core *Paramoeba/Neoparamoeba* clade remains unresolved, as neither their inclusion nor alternative positions can be statistically rejected (Table S4). Resolving the relationships within the *Paramoeba/Neoparamoeba* clade will require expanded taxon and gene sampling.

The phylogenetic position of the new isolate varies across different analyses (Table S3). In 18S gene-based phylogenies, it alternates between forming a sister-group relationship with *P. eilhardi* or with *P. aparasomata*+*P. karteshi*, with both placements remaining statistically unresolved (Table S3). The interspecific divergence among COI gene sequences within the EDAK clade is relatively low (12.5%–14.5%), making it challenging to obtain strong support in COI gene phylogenies (Figure 6). However, multiple lines of evidence, including morphological characteristics, COI phylogeny, and some statistical support (Table S4), favor a closer relationship between the new isolate and *P. karteshi*+*P. aparasomata*. Given its morphological similarities (e.g., presence of PLO and other morphological features), the new isolate is likely a sister species to *P. karteshi*. However, further data are needed to confirm these relationships.

### Endosymbiont phylogeny using raw nanopore metabarcoding data

We constructed an 18S gene phylogeny of the endosymbiont using error-uncorrected nanopore data (Figure S3). The 18S gene sequence from the new isolate was obtained through co-amplification with host DNA and sequenced using Oxford Nanopore technology. Nanopore sequencing has been widely applied in metabarcoding studies across various lineages (Chang et al. 2024, Stoeck et al. 2024). Although recent improvements in nanopore technology have reduced error rates, our raw data still contained a number of indels and substitutional errors (data not shown). However, with bioinformatics filtering and consensus-building, high-quality data can be obtained when sequencing depth is sufficient, as demonstrated in this study for the host 18S gene data (Figure 5; (Chang et al. 2024)). Using unpolished data, we reconstructed a PLO 18S gene phylogeny congruent with previously published phylogenies (Sibbald et al. 2017, Volkova et al. 2019, Young et al. 2014), reaffirming the co-evolutionary relationship between the host and its endosymbiont. This approach is promising for amoeboid metabarcoding due to its efficiency, as it eliminates the need for labor-intensive cloning to capture intrastrain variation and offers a faster turnaround time compared to traditional sequencing techniques.

### Observation on PLO morphology, movement and association with microtubules

The endosymbiotic relationship of the PLO and *Paramoeba/Neoparamoeba* is a classic and intriguing relationship of eukaryote-to-eukaryote symbiosis that is not dependent on photosynthesis. This observation has led much research to study this relationship at cellular and genomic levels with an evolutionary context (Tanifuji et al. 2017, Dyková et al. 2003, David et al. 2015). Studies have shown that the PLO is an aflagellate, obligatory intracellular endosymbiont surrounded by a single membrane, not derived from its host and resides near the host nucleus. It is often binuclear, with nuclei positioned at opposite poles, and its cell body is predominantly occupied by a large mitochondrion structurally similar to other kinetoplastids.

PLO possesses a highly reduced genome and is closely related to parasitic protists within Kinetoplastida (Dyková et al. 2003, Tanifuji et al. 2011). However, a lot remain to be unraveled about its biology and role in survival or pathogenicity of the host amoeba. While it is not our intention to describe a detailed relationship between the PLO and host amoeba, it this study we present a few important observations based on the new isolate that can contribute towards understanding this intimate association.

PLOs are frequently observed near the nucleus, sometimes leaving a pit-like, round-shaped compression mark on the host nucleus (Dyková et al. 2003). In the newly isolated strain, the PLO is consistently in direct contact with the host nucleus (Figures 2, 3). While PLO morphology is generally similar across observations, notable variations have been reported, including differences in the middle region’s morphology, PLO size and shape, and the intracellular arrangement of the nucleus and other components (Volkova et al. 2019). One particularly interesting observation from our staining experiments using DAPI and CellMask is that neither the PLO nucleus nor its surrounding membrane was stained (Figure 3). However, other studies have reported strong DAPI staining within the PLO (Volkova and Kudryavtsev 2017, Dyková et al. 2003). All these recorded variations are intriguing and might have taxonomic implications (see Volkova et al. 2019).

The visibility of PLO is more apparent and prominent in ICC staining of microtubules (Figures 3, S1). Each PLO is surrounded by single or bundles of microtubules (Figures 3). The anti-tubulin antibodies used to stain the area surrounding the PLO appear to react more strongly than those targeting the host amoeba’s MT (Figures 3, S1). Even under conditions where the host amoeba’s MT are not well preserved, the MT encircling the PLO remain consistently visible with robust signal (Figures 3A).

Despite multiple staining attempts, it remains unclear whether the microtubules (MT) surrounding the PLO are a continuation of the host’s MT or originate from the endosymbiont itself. Genome analysis of PLO has revealed a highly reduced genome, including the loss of most genes associated with flagellar structures. Additionally, the main genetic subunits of MT, α- and β-tubulins, have undergone mutations and motif losses that could potentially hinder their structure and function (Tanifuji et al. 2011). However, previous ultrastructural studies have identified MT-like structures within the PLO (Perkins and Castagna 1971, Dyková et al. 2003).

Some of these structures were linked to the division of the endosymbiont, while others—larger than typical MT—were reported to run along the middle part of the PLO cell (Perkins and Castagna 1971). Although this ultrastructural study did not capture the entire MT network along the middle part of the PLO cell, both the thickness (interpreted here as bundles) and the length of these MT structures resemble the strongly stained MT observed around the PLO in our study (Figures 3, S1). The study by (Dyková et al. 2003) using high-quality TEM sections clearly demonstrated the presence of MT running along thin membrane section in the middle part of the PLO cell. Based on these observations and our findings, it is likely that PLO produces its own cytoplasmic MT as part of its cellular structure. These MT structures are present in every examined PLO (Figures 3, S1) and appear to be permanent components, likely playing structural and functional roles that remain to be elucidated. The discovery of a microtubular system in PLO is significant, given the loss of flagellum and the genetic modifications related to cytoskeleton reported in its genome (Tanifuji et al. 2011). Further investigation is underway to determine the origin and function of the MT network and its role in cell structure and interaction with the host amoeba.

In a related context, it is intriguing to consider whether the microtubular system of the host or the PLO have any role in maintaining the close proximity between the host nucleus and the PLO. Observations of PLO movement indicate that during cytoplasmic streaming, the PLO moves around the nucleus (Data 1 timelapse). Although the PLO was seen moving relatively freely around the nucleus, it was never observed to complete a full circle around it. This restricted movement may be due to its association with the host MT, although this was not clearly evident in our staining. It is important to investigate how the MT system in the PLO interacts and functions with the host amoeba. While these observations provide valuable insights, much remains to be uncovered regarding PLO movement within the host cell and its interaction with the host nucleus.

**Taxonomic Appendix** Based on Adl et al. (2019): Amorphea: Amoebozoa: Discosea: Flabellinia: Dactylopodida: *Paramoeba* Schaudinn, 1896, *sensu* Volkova et al. 2019

***Paramoeba daytoni* n. sp. Tekle 2025**

**Diagnosis:** Locomotive forms predominantly flattened and triangular, with dactylopodial subpseudopodia extending dynamically. Length of locomotive form 31.15–78.43 µm (average 54.73 µm), breadth 9.23–54.53 µm (average 23.85 µm). Subpseudopodia 3.55–31.34 µm long (average 9.81 µm), typically 7–10 per cell. Locomotion speed 13–19.69 µm/min (average 15.7 µm/min). Single vesicular nucleus 5.66–9.99 µm in diameter (average 7.75 µm). Usually more than one PLOs, occasionally up to four; PLOs 2.02–4.67 µm long (average 3.07 µm) and 1.76–3.19 µm broad (average 2.53 µm), typically bean- or round-shaped.

**Type locality:** Marine, seaweed from a public beach of Daytona Beach, Northeastern Coastal regions of Florida (66.3369944N, 33.6598806E).

**Type material and molecular data:** A type culture will be kept in Tekle laboratory cryotank storage; GenBank accession numbers XXXXX-XXXXXX (SSU-rDNA, 18S, gene), XXXXXX (COI gene).

**Etymology:** This species is named after the type locality of “Daytona” Beach, Florida where the new isolate was discovered.

## ACKNOWLEDGEMENTS

This work is supported by the National Science Foundation EiR (2401946) and the National Institutes of Health (1R15GM116103-02) to YIT. We would like to thank Dr. Joseph Ryan and Dr. Yuriy Bobkov for organizing our visit to the Whitney Laboratory for Marine Bioscience, Florida.

## SUPPORTING INFORMATION

**Figure S1.**
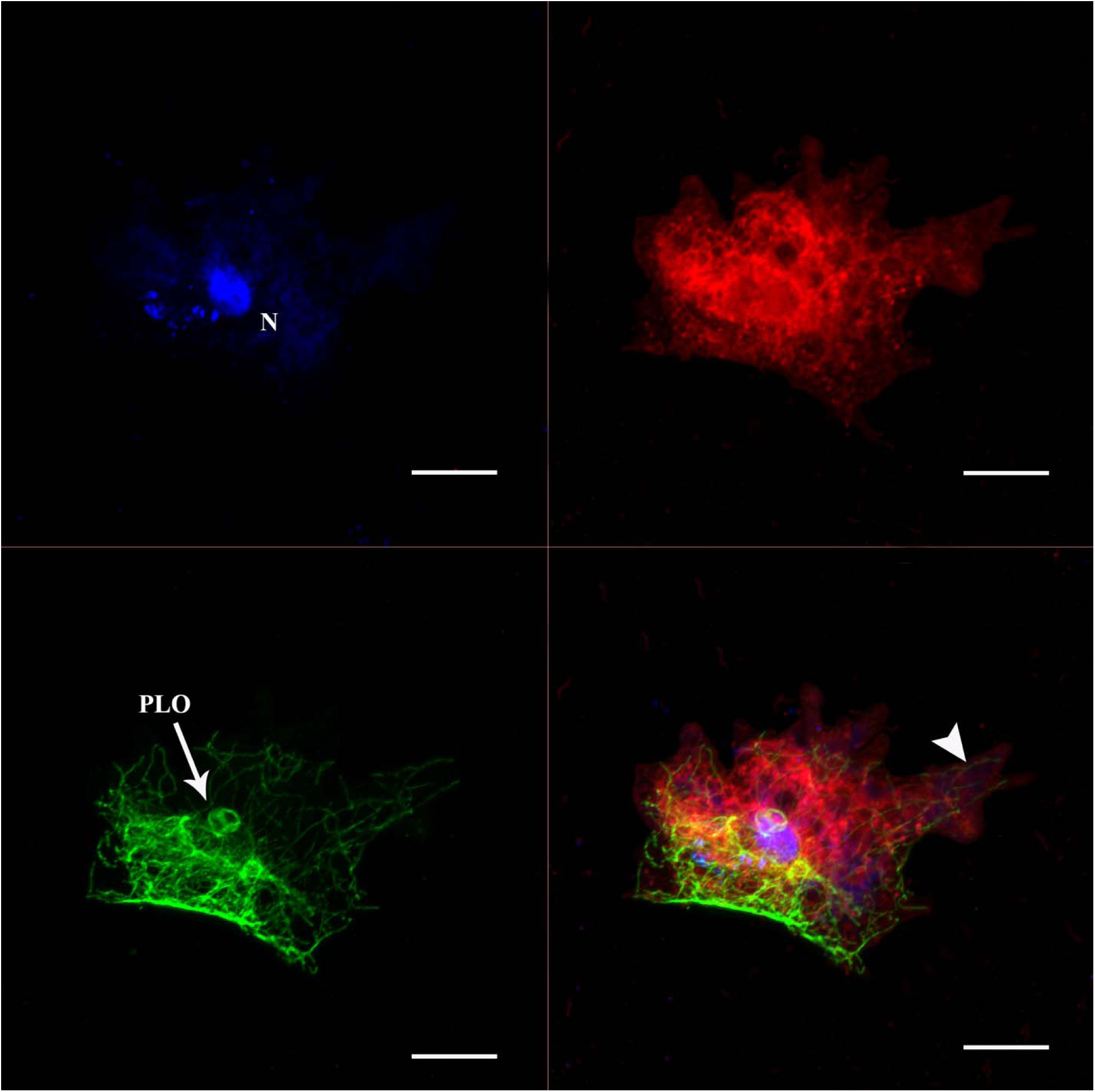
Confocal maximum intensity projection split images of triple staining in *Paramoeba daytoni* n. sp., showing DNA (blue, A), microtubules (green, B), plasma membrane (red, C), and a merged image of all three (D). The staining reveals a well-preserved network of host amoeba microtubules (MTs) along with two PLO-associated MTs (C, D). Notably, the host MTs surrounding the nucleus form a network that appears to support the host nucleus, with a weak connection observed between these MTs and the PLO MTs (arrowhead, C). The subpseudopodia are supported by a few extending MTs (arrow, D).

**Figure S2.**
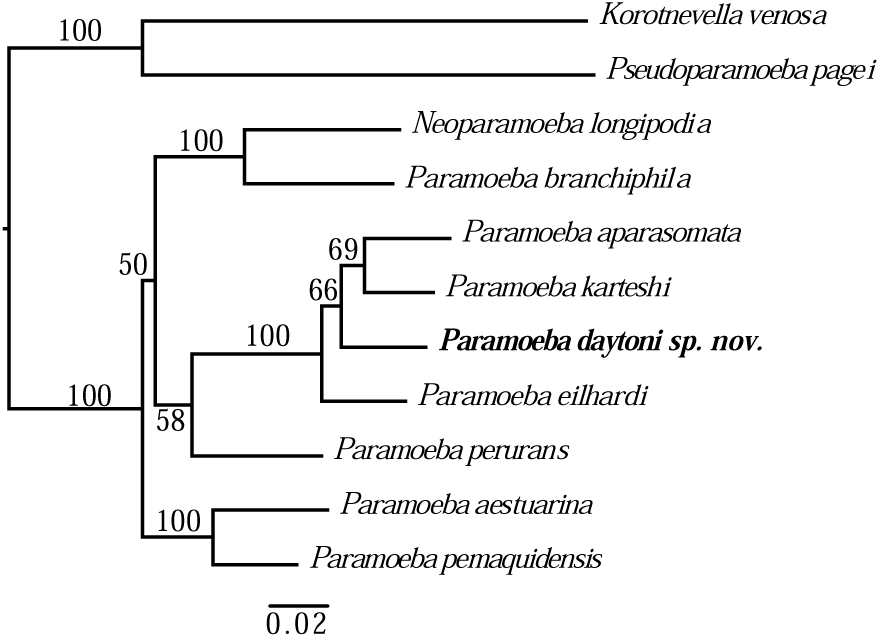
IQ-TREE maximum likelihood phylogeny based on nine ingroup taxa combining 18S and COI genes of *Paramoeba/Neoparamoeba* clade species, highlighting the position of *Paramoeba daytoni* n. sp. The alignment, generated using AliView, includes 2,699 nucleotide positions. Numbers at nodes represent IQ-TREE bootstrap values. Branches are drawn to scale.

**Figure S3.**
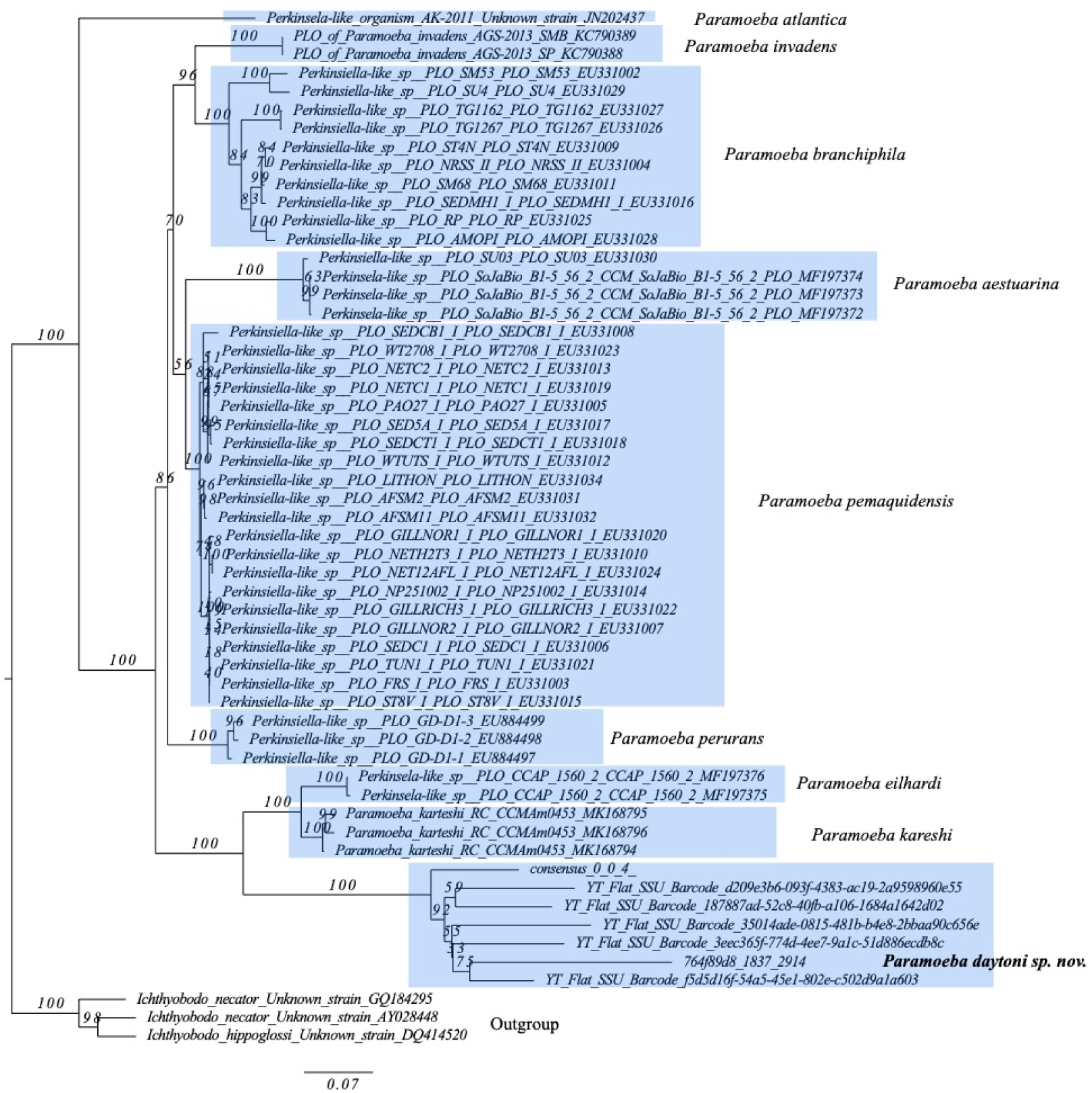
IQ-TREE maximum likelihood phylogeny based on PLO 18S gene of *Paramoeba/Neoparamoeba* clade species, highlighting the position of the PLO of *Paramoeba daytoni* n. sp. Numbers at nodes represent IQ-TREE bootstrap values. Branches are drawn to scale.

**Data 1:** Time-lapse movie tracking the movement of the PLO within several host amoeba cells using DIC imaging. The PLOs are observed migrating (arrow) along the periphery of the host nucleus but do not complete a full circular rotation.

### Supplementary Table

**Table S1.** SSU-rDNA (18S) Intrastrain Variation of *Paramoeba/ Neoparamoeba* species.

| Species | Minimum | Maximum | Average |
| --- | --- | --- | --- |
| *Paramoeba invadens* | 0.08% | 2.68% | 1.16% |
| *Paramoeba branchiphila* | 0.80% | 4.97% | 2.61% |
| *Neoparamoeba longipodia* | 0.05% | 1.33% | 0.98% |
| *Paramoeba perurans* | 0.00% | 1.67% | 0.81% |
| *Paramoeba eilhardi* | 1.76% | 2.54% | 2.12% |
| *Paramoeba karteshi* | 1.21% | 1.64% | 1.38% |
| *Paramoeba daytoni* | 0.00% | 1.71% | 0.94% |
| *Paramoeba aparasomata* | 0.49% | 2.31% | 1.79% |
| *Paramoeba pemaquidensis* | 0.00% | 3.58% | 2.03% |
| *Paramoeba aestuarina* | 0.14% | 5.54% | 3.26% |

**Table S2.**
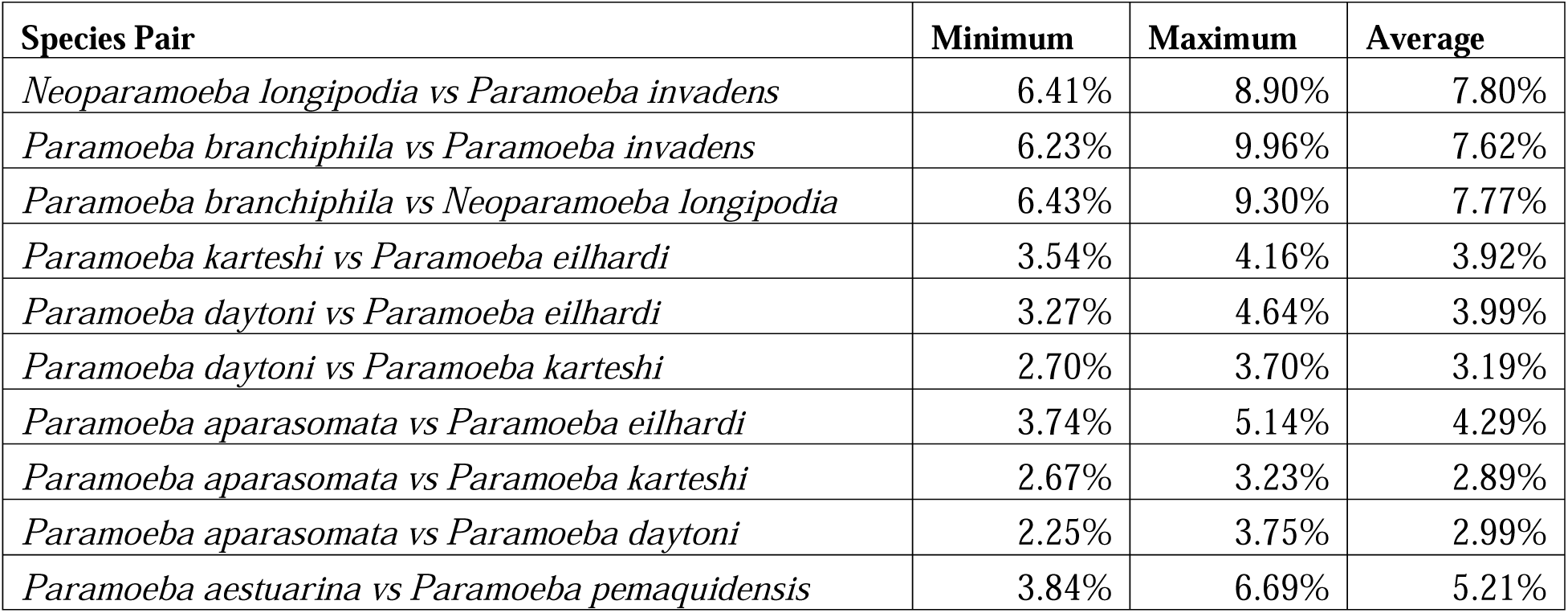
SSU-rDNA (18S) interspecific variation of *Paramoeba/ Neoparamoeba* clades.

| Species Pair | Minimum | Maximum | Average |
| --- | --- | --- | --- |
| *Neoparamoeba longipodia* vs *Paramoeba invadens* | 6.41% | 8.90% | 7.80% |
| *Paramoeba branchiphila* vs *Paramoeba invadens* | 6.23% | 9.96% | 7.62% |
| *Paramoeba branchiphila* vs *Neoparamoeba longipodia* | 6.43% | 9.30% | 7.77% |
| *Paramoeba karteshi* vs *Paramoeba eilhardi* | 3.54% | 4.16% | 3.92% |
| *Paramoeba daytoni* vs *Paramoeba eilhardi* | 3.27% | 4.64% | 3.99% |
| *Paramoeba daytoni* vs *Paramoeba karteshi* | 2.70% | 3.70% | 3.19% |
| *Paramoeba aparasomata* vs *Paramoeba eilhardi* | 3.74% | 5.14% | 4.29% |
| *Paramoeba aparasomata* vs *Paramoeba karteshi* | 2.67% | 3.23% | 2.89% |
| *Paramoeba aparasomata* vs *Paramoeba daytoni* | 2.25% | 3.75% | 2.99% |
| *Paramoeba aestuarina* vs *Paramoeba pemaquidensis* | 3.84% | 6.69% | 5.21% |

**Table S3.** Clade Support and Recovery Across Datasets Aligned and Analyzed Using Different Phylogenetic Algorithms.

| Data analyzed | IBL | PA | E-DAK | ED-AK | EDAK+PA | Pe+IBL | EDAK+IBL | Pe+IBL | Pe+(PA+EDAK) | Pe+EDAK |
| --- | --- | --- | --- | --- | --- | --- | --- | --- | --- | --- |
| IQ-Aliview_2300 | + | + | + | - | - | + | + | - | - | - |
| IQ-Mafft_2310 | + | + | + | - | - | + | + | - | - | - |
| IQ-Aliview_1750 | + | + | - | + | + | - | - | + | - | - |
| IQ-Mafft_1760 | + | + | - | + | + | - | - | + | - | - |
| RaxML-Aliview_2300 | + | + | + | - | + | - | - | - | + | - |
| RaxML-Mafft_2310 | + | + | - | + | - | + | + | - | - | - |
| RaxML-Aliview_1750 | + | + | + | - | + | - | - | + | - | - |
| RaxML-Mafft_1760 | + | + | - | + | - | + | + | - | - | - |
| MrBayes-Aliview_2300 | + | + | + | - | - | - | - | - | - | + |
| MrBayes-Mafft_2310 | + | + | + | - | - | + | - | - | - | - |
| MrBayes-Aliview_1750 | + | + | - | + | + | - | - | - | + | - |
| MrBayes-Mafft_1760 | + | + | - | + | + | - | - | - | + | - |
| Total recovered | 12 | 12 | 6 | 6 | 6 | 5 | 4 | 3 | 3 | 1 |

Taxon and clade name key: **IBL** - (*P. invadens* + *P. branchiphila*) + *Neoparamoeba longipodia*; **PA** - *P. pemaquidensis* + *P. aestuarina*; **Pe** - *P. perurans*; **E** - *P. eilhardi*; **D** - *P. Dayton sp. n.*; **A** - *P. aparasomata*; **K** - *P. kareshi*.

**Table S4.** Approximately Unbiased (AU) Test for Alternative Topologies and Taxon Placement in Phylogenetic Analyses.

| Tree | p-AU |
| --- | --- |
| **Hypothesis I** |  |
| Main Tree Fig. 5. ((IBL,(Pe,(PA,(E,(D,(A,K)))))) | 0.4 |
| Alternative: ((IBL,(Pe,(PA,((E,D),(A,K)))))) | 0.3 |
| **Hypotheses II** |  |
| Main Tree Fig. 5. ((IBL,(Pe,(PA,(E,(D,(A,K)))))) | 0.035* |
| Alternative: (((PA,(E,(D,(A,K))))),IBL), Pe) | 0.241 |
| Alternative: (((PA,(E,(D,(A,K))))),Pe), IBL) | 0.877 |

\*p<0.05 is rejected.
Taxon and clade name key: **IBL** - (*P. invadens* + *P. branchiphila*) + *Neoparamoeba longipodia*; **PA** - *P. pemaquidensis* + *P. aestuarina*; **Pe** - *P. perurans*; **E** - *P. eilhardi*; **D** - *P. Dayton sp. n.*; **A** - *P. aparasomata*; **K** - *P. kareshi*.

**DATA S1 timelapse – available at this YouTube link.** https://youtu.be/Psd-cwIQAjY

